# Remote Ischemic Preconditioning Ameliorates Anthracycline-induced Cardiotoxicity and Preserves Mitochondrial Integrity

**DOI:** 10.1101/2020.03.20.000729

**Authors:** Carlos Galán-Arriola, Rocio Villena-Gutiérrez, María I Higuero-Verdejo, Iván A. Díaz-Rengifo, Gonzalo Pizarro, Gonzalo J. López, Antonio de Molina-Iracheta, Claudia Pérez-Martínez, Rodrigo D. García, David González-Calle, Manuel Lobo, Pedro L Sánchez, Eduardo Oliver, Raúl Córdoba, Valentin Fuster, Javier Sánchez-González, Borja Ibanez

## Abstract

**Aims:** Anthracycline-induced cardiotoxicity (AIC) is a serious adverse effect in a significant proportion of cancer patients. A central mechanism of AIC is irreversible mitochondrial damage. Despite major efforts, there are currently no effective therapies able to prevent AIC.

**Methods and Results:** Forty Large-White pigs were included. In Study 1, 20 pigs were randomized 1:1 to remote ischemic pre-conditioning (RIPC, 3 cycles of 5 min leg ischemia followed by 5 min reperfusion) or no pretreatment. RIPC was performed immediately before each of five intracoronary doxorubicin injections (0.45 mg/kg) given at weeks 0, 2, 4, 6, and 8. A group of 10 pigs with no exposure to doxorubicin served as healthy controls. Pigs underwent serial cardiac magnetic resonance (CMR) exams at baseline and at weeks 6, 8, 12, and 16. After 16-week CMR, pigs were sacrificed and tissue samples collected. In study 2, 10 new pigs received 3 doxorubicin injections (with/out preceding RIPC) and were sacrificed 2 weeks after the third dose.

In Study 1, LVEF remained unchanged in doxorubicin-treated pigs until week 6 (time of the fourth doxorubicin injection). From there on, LVEF progressively declined, but LVEF depression was blunted animals receiving RIPC before doxorubicin (RIPC-Doxo), which had a significantly higher LVEF at week 16 than doxorubicin treated pigs that received no pretreatment (Untreated-Doxo) (41.5±9.1% vs 32.5±8.7%, p=0.04). Preserved LVEF was mainly due to conserved contractile function, as evidenced by smaller LVESV, and better regional contractile function. In Study 2, transmission electron microscopy (TEM) after 3 doxorubicin doses showed fragmented mitochondria with severe morphological abnormalities in RIPC+Doxo pigs, together with upregulation of fission proteins and autophagy markers on western blot. At the end of the 16-week Study 1 protocol, TEM revealed overt mitochondrial fragmentation with structural fragmentation in Untreated-Doxo pigs, whereas interstitial fibrosis was significantly less severe in the RIPC+Doxo pigs.

**Conclusion:** In a translatable large animal model of AIC, RIPC applied immediately before each doxorubicin injection resulted in preserved cardiac contractility with significantly higher long-term LVEF and less cardiac fibrosis. RIPC prevented mitochondrial fragmentation and dysregulated autophagy from the early stages of AIC. RIPC is a promising intervention for testing in clinical trials in AIC.

## Introduction

Anthracyclines, used alone or in combination with other approaches, remain the first line therapy for many forms of cancer. Anthracycline-induced cardiotoxicity (AIC) is a well-known side effect of these agents, limiting the total lifetime cumulative dose a patient can receive^1^. The mechanisms by which anthracyclines generate cardiotoxicity seem to converge on damage to mitochondria^2^.

Several definitions of cardiotoxicity have been proposed, with a consensus conception of cancer–therapy-related cardiac dysfunction as a reduction in left ventricular ejection fraction (LVEF) of 10 absolute points to a value below the normal range for the imaging technique^3–6^. Severe AIC is associated with chronic heart failure (HF), which translates into increased mortality^6^.

Current treatment strategies for established AIC include standard HF therapies (beta-blockers and ACEs inhibitors, among others). Early intervention with these therapies is of limited effect, providing partial recovery of cardiac function in only some patients^7^. With the exception of dexrazoxane, which reduced cardiac injury in a pediatric population^8^, specific therapies able to prevent the development of AIC are lacking^8^. Given the central role of mitochondria in AIC, several interventions targeting mitochondrial processes have been tested, including antioxidants, iron chelators, and cell therapy; however, despite promising preclinical results, these approaches have failed to demonstrate a clinical benefit^9^.

Remote ischemic conditioning is the process whereby brief, reversible episodes of ischemia and reperfusion in one vascular bed (e.g. an arm) confers a global protection that renders remote tissues and organs resistant to injury^10^. A central mechanism underlying the benefits of remote ischemic conditioning is mitochondrial protection^10^. Conditioning before the index injury (remote ischemic PREconditioning (RIPC)^11^ has been shown to be very effective; however, initiation of conditioning during the index injury (remote ischemic PERconditioning) confers a weaker protection^12^. Nevertheless, anthracycline administration is a fully controlled and programmed intervention, and the cardiac injury event is thus predictable. Therefore, given the central role of mitochondrial damage in AIC, RIPC seems a good candidate therapy for testing in this context. In this study, we tested RIPC as a cardioprotective strategy in a validated large animal model of AIC^13^. Monitoring by cardiac magnetic resonance (CMR) revealed that the severe cardiac dysfunction associated with serial intracoronary doxorubicin administration is significantly attenuated by RIPC applied immediately before each anthracycline administration. RIPC prevented severe anthracycline-induced mitochondrial fragmentation and dysregulated autophagy.

## Methods

### Study design

All animal studies were conducted at the CNIC and approved by the local CNIC Institutional and regional animal research committees. All animal procedures conformed to EU Directive 2010/63EU and Recommendation 2007/526/EC regarding the protection of animals used for experimental and other scientific purposes.

The study design is summarized in **Figure 1**. In Study 1, we tested the efficacy of RIPC in a model of overt AIC with long-term evaluation: 20 Large-White male pigs (weight ∼30 kg) received a series of 5 intracoronary doxorubicin injections in order to induce severe AIC^13^. The 0.45 mg/kg injections were spaced 2 weeks apart (weeks 0, 2, 4, 6, and 8). Before commencing injections, animals were randomized 1:1 to RIPC or no pretreatment. There were thus two AIC treatment groups: those with preconditioning (RIPC-Doxo) and those with no pretreatment (Untreated-Doxo). RIPC consisted on 3 cycles of 5 minutes lower limb ischemia followed by 5 minutes reperfusion before each doxorubicin injection. All pigs underwent serial CMR exams at weeks 0, 6, 8, 12, and 16. After the 16-week CMR exam, animals were sacrificed, and cardiac tissue was harvested for histology, transmission electron microscopy (TEM), and protein expression analysis. Another group of 10 pigs with no exposure to doxorubicin served as healthy controls.

**Figure 1:**
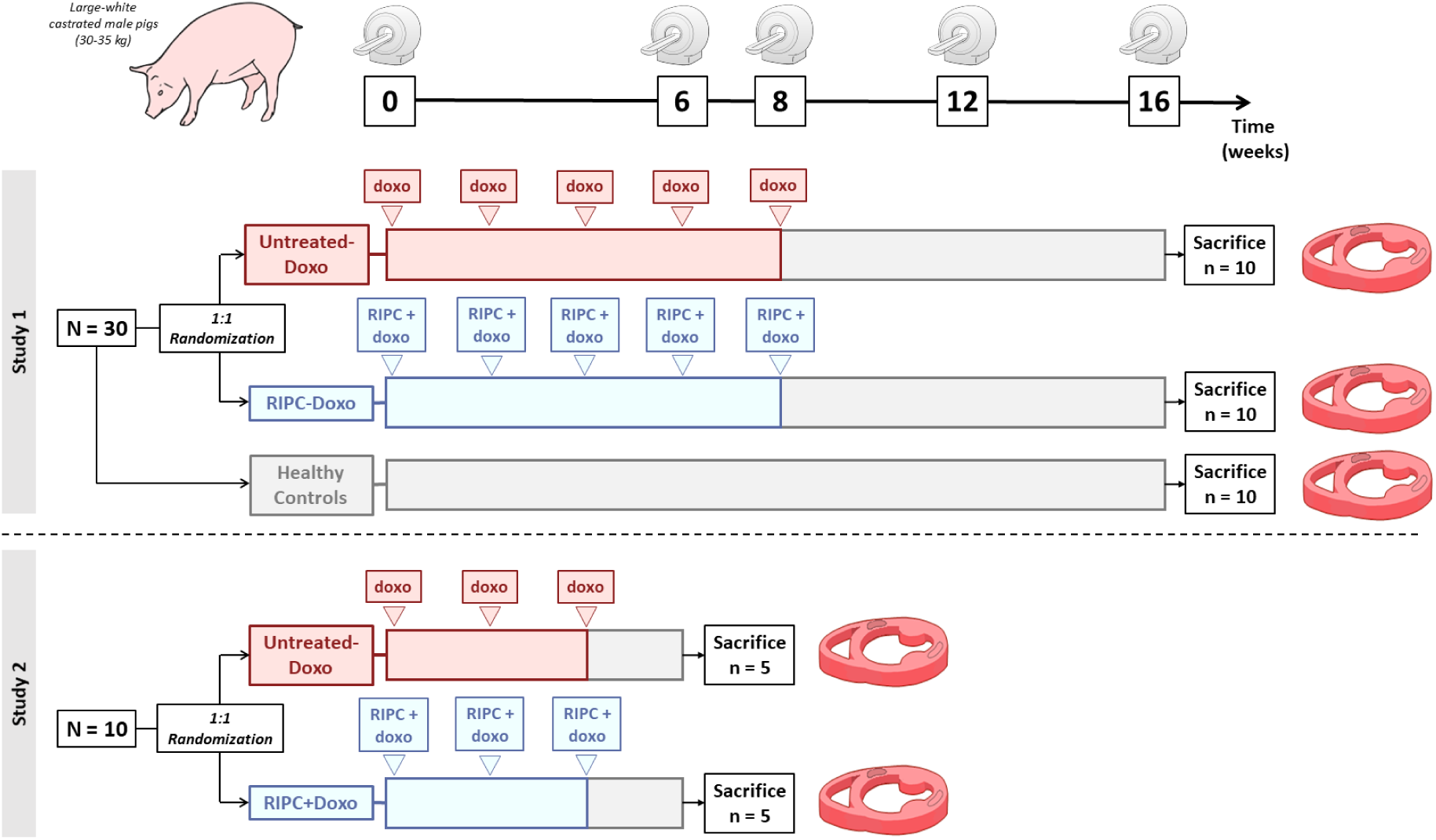
Study design. Study 1 consisted of 30 castrated Large White male pigs; 20 pigs were randomized the AIC protocol with no pretreatment (Untreated-Doxo) or preceded at each cycle by RIPC (RIPC-Doxo). An additional 10 animals were monitored as healthy controls. CMR scans were performed at baseline (week 0) and at 6, 8, 12, and 16 weeks. Animals were sacrificed, and hearts were excised for ex-vivo analysis. Study 2 consisted of 10 animals randomized to Untreated-Doxo or RIPC-Doxo and sacrificed 2 weeks after the third doxorubicin dose. CMR scans were performed at baseline (week 0) and at 6 weeks.

In Study 2, 10 new pigs (randomized to RIPC or no treatment) were administrated with only 3 doxorubicin intracoronary injections, a “subclinical” protocol associated with preserved cardiac function^13^. The RIPC-Doxo and Untreated-Doxo pigs in Study 2 underwent CMR exams at weeks 0 and 6. After the 6-week CMR exam, pigs were sacrificed and cardiac tissue harvested for histology, TEM, and protein expression analysis.

### Remote ischemic preconditioning

RIPC was induced by securing a tourniquet around the right hind leg to produce transient ischemia. Total ischemia was assured by checking visually for the congestive reaction in the lower extremity and by doppler echography, pressing the tourniquet until artery flow ceased. Each of the three RIPC cycles consisted of 5 minutes of total ischemia followed by 5 minutes of reperfusion.

### Intracoronary doxorubicin injection

Doxorubicin was administered according to a previously detailed protocol^13^. In brief, animals were anesthetized and endotracheally intubated. The femoral artery was then accessed by the Seldinger technique, and a 5F sheath was inserted. Pigs were anticoagulated with 150 IU/kg of intravenous heparin, and a 4F coronary diagnostic catheter was inserted via a femoral sheath and placed at the origin of the left coronary artery. Under angiography guidance, a 0.014 mm coronary guide wire was positioned distally in the left anterior descending (LAD) coronary artery. The catheter was docked selectively in the proximal LAD, and a 0.45 mg/kg dose of doxorubicin (Farmiblastina®, Pfizer) diluted in 30 ml saline was given as a slow bolus injection over 3 minutes. Electrocardiographic and hemodynamic parameters were monitored during doxorubicin administration. Once infusion was complete, normal coronary flow was documented by coronary angiography, the material was removed, and the animal was allowed to recover.

### Cardiac magnetic resonance imaging

All studies were performed with a Philips 3-T Achieva Tx whole body scanner (Philips Healthcare, Best, The Netherlands) equipped with a 32-element phased-array cardiac coil. The CMR protocol included a standard segmented cine steady-state free-precession (SSFP) sequence to provide high-quality anatomical references, T1-mapping, and T1W late gadolinium enhancement (LGE) sequences. The imaging parameters for the cine SSFP sequence were as follows: field of view (FOV), 280 × 280 mm; slice thickness, 6 mm with no gaps; repetition time (TR), 2.8 ms, echo time (TE), 1.4 ms; flip angle, 45°; cardiac phases, 30; voxel size, 1.8 × 1.8 mm; and number of excitations (NEX), 3. The T1-mapping sequence (Modified Look-Locker Inversion recovery [MOLLI]) was based on a 5(3)3 scheme using a single-shot steady-state free precession readout sequence (TR/TE/Flip angle= 2.1 ms/1.05 ms/350) with an in-plane acquisition resolution of 1.5×1.8 mm^2^ and an 8 mm slice thickness. The T1 mapping sequence was triggered at mid-diastole and acquired from a single short-axis mid-apical slice. LGE imaging was performed 15 min after intravenous administration of 0.2 mM/kg gadopentetate dimeglumine contrast agent using an 3D inversion-recovery spoiled turbo field echo sequence (TR/TE/Flip angle= 2.4ms/1.13ms/100) with an isotropic resolution of 1.5×1.5×1.5 mm^3^ on a FOV of 340×340×320 mm^3^ in the FH, LR, and AP directions. Data were acquired in mid-diastole with a 151.2 ms acquisition window. Acquisition was accelerated using a net SENSE factor of 2.25 (1.5×1.5 in the AP and LR directions) with a bandwidth of 853Hz per pixel. Inversion time was adjusted before acquisition using a look-locker scout sequence with different inversion times to ensure proper nulling of the healthy myocardium signal. For the analysis, 3D volume was reconstructed in short axis, 2 chamber, and 4 chamber views with a slice thickness of 6mm.

### Image analysis

CMR studies were analyzed by an experienced observer and reviewed by a blinded experienced independent observer using dedicated software (IntelliSpace Portal, Philips Healthcare, Best, the Netherlands). LV cardiac borders were traced in each cine image to obtain LV end-diastolic mass, LV end-diastolic volume (LVEDV), end-systolic volume (LVESV), and Left LV ejection fraction (LVEF). Wall thickening values for the assessment of infused (LV region supplied by the LAD) and remote contractility were also obtained by tracing cardiac borders in systole and diastole. T1 maps were generated by using a maximum-likelihood expectation-maximization algorithm and fitting the MR signal to a T1 inversion recovery with 3 independent model parameters. T1 relaxation maps were quantitatively analyzed by placing a wide transmural region of interest at the infused myocardial area (irrigated by the LAD) of the corresponding slice in all studies.

### Ex-vivo analysis

After sacrifice, samples of the doxorubicin-infused region (anterior wall) and the remote area (posterior wall) were collected for histology, TEM, and protein expression analysis. For histology, samples were fixed in 4% formalin and embedded in paraffin, and 4-μm sections were cut and stained with hematoxylin and eosin, Masson trichrome, and Sirius Red. Sirius Red sections were scanned, and 20 × 20 magnification images were captured for collagen quantification using a modified macro^14^.

### Transmission electron microcopy

Samples from the infused area were maintained in 4% glutaraldehyde in 10% paraformaldehyde for 24-48 hours. Tissues were then post-fixed in 1% osmium tetroxide in water for 1 hour at room temperature. Samples were washed with water and block-stained with 0.5% uracil acetate in water for 10 minutes. Samples were then dehydrated through a series of aqueous alcohol solutions (30%, 50%, 70%, 95%, and 100%) and a final passage through acetone. After this, samples were included in Durcupan epoxy resin through increasing resin:acetone mixtures (1:3 then 3:1) and, finally, pure resin. Resin-included samples were polymerized in an oven at 60°C for 48 hours. Ultrathin (60 nm) slices were cut with a Leica Ultracut S ultramicrotome and were deposited on 200 mesh copper grids. Grids were counterstained with uranyl acetate and lead citrate. Images were obtained with a Jeol Jem1010 (100 KV) transmission electron microscope linked to a Gatan camera (Orius 200 SC model); acquired images were processed with Digital Micrograph software. Individual images were acquired using ImageJ software (1.52p Wayne Rasband from the US National Institutes of Health) at the following magnifications: 6000for X tissue evaluation, 10000X for mitochondrial quantification, and 40000X for the assessment of mitochondrial cristae.

### Western blotting

Heart tissue from infused area was lysed in RIPA buffer supplemented with a protease and phosphatase inhibitor cocktail. Protein content was quantified with the Bio-Rad BCA protein assay. Protein samples were separated by SDS–polyacrylamide gel electrophoresis, and proteins were transferred to nitrocellulose membranes. After blocking, membranes were incubated with the following antibodies: BECLIN1 (Cell Signaling, 3738S), p62 (Cell Signaling, 5114S), DRP1 (Cell Signaling, 8570S), and GAPDH (Abcam, ab8245) at 4 °C overnight. Bound antibodies were detected after staining with a corresponding secondary antibody. Quantitative densitometric analysis was performed using ImageJ Fiji software.

### mtDNA quantification

mtDNA was quantified by quantitative real-time PCR using SYBR Green, and mtDNA content was presented as the ratio of mitochondria-encoded 16S to nuclear-encoded HPRT. 16S primers: forward 5′-CGATGTTGGATCAGGACACC-3′, reverse 5′-CTGAGACGCGTTTGTGAAGTT-3′. HPRT primers: forward 5′-GGCCAGTTCGGGAATGATCT-3′, reverse 5′-CCCCCAGTCCCCCAAATCTA-3′.

### Statistical analysis

According to the data distribution, continuous variables were calculated as mean±SD or median ± IQR. Data normality was assessed with the Saphiro-Wilk test. Cardiac mass and volume data were indexed by body weight using the modified Brody formula. Differences were considered statistically significant at p<0.05. All data were analyzed with RStudio (RStudio Team (2015): Integrated Development for RStudio, Inc., Boston, MA), and graphics were created with ggplot2.

Sample size in each treatment group (RIPC-Doxo and Untreated-Doxo) was calculated in order to detect an 8% between-group difference in LVEF at 16-week CMR (study primary endpoint) with an SD of 6.5%, no casualties (previously defined in the animal model^13^), a power of 0.8, and 0.05 of bilateral significance. The primary endpoint is similar to that in other cardioprotection studies, and the sample size calculation resulted in 10 animals per group. Two-group comparisons were by *t*-test, taking account of RIPC exposure at the end of the study (16 weeks). Comparisons over time within each group were made by repeated ANOVA measurements with Bonferroni correction.

## Results

During doxorubicin injections, none of the pigs showed adverse events (changes in ECG or systolic arterial pressure). One pig randomized to RIPC died immediately after the third anthracycline injection due to a complication related to the invasive procedure and was replaced to maintain the pre-specified sample size. Another pig from Untreated-Doxo group died after the final CMR exam at 16 weeks, we decided not to use ex-vivo samples for water quantification and TEM/WB from this animal.

### Remote ischemic preconditioning prevents anthracycline-induced cardiac dysfunction

The series of 5 doxorubicin injections in Study 1 caused a progressive deterioration of LV systolic function in pigs allocated to the non-RIPC AIC group (Untreated-Doxo). LVEF remained unchanged until the fourth doxorubicin injection. From there on, LVEF progressively deteriorated to the end of the study. LV systolic function deterioration was significantly blunted in pigs receiving RIPC before each doxorubicin injection (RIPC-Doxo group) (**Figure 1**). LVEF at week 16 (primary study endpoint) was significantly higher in the RIPC-Doxo group (41.5 % ± 9.1 and 32.5 ± 8.7 for RIPC-Doxo vs Untreated-Doxo; p = 0.04). Cardiac dysfunction was already overt at week 12 CMR (LVEF = 43.2 % ± 8.1 and 33.4 ± 9.1 in RIPC-Doxo vs Untreated-Doxo; p = 0.02) (**Figure 2**.**A**). There was no between-group difference in 16-week LVEDV (156.0 ml ± 38.4 and 168.0 ± 69.7 in RIPC-Doxo vs Untreated-Doxo; p = 0.63), whereas LVESV was non-significantly lower in the RIPC group (93.1 ml ± 35.2 and 118.0 ± 64.3 in RIPC-Doxo vs Untreated-Doxo; p = 0.30) (*Supplementary material online, Figure 1*). All CMR data from Study 1 are detailed in *Supplementary material online, Table 1*.

**Figure 2:**
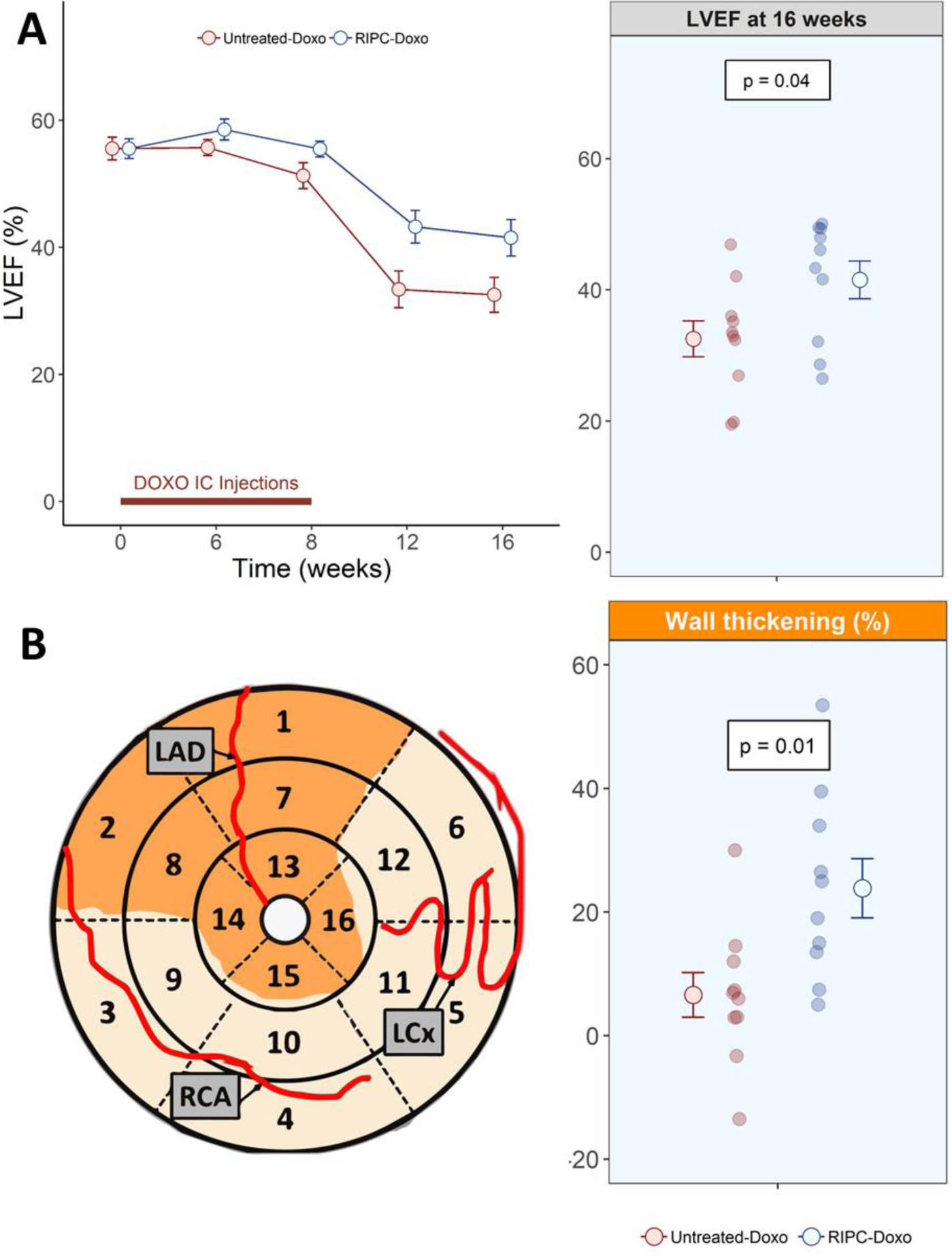
Functional CMR in Study 1. (**A**) LVEF over time (left) and at 16 weeks (right). Grouped data are presented as mean±SE and individual data at the end of the study as a dotplot. N=10 animals per group. The significance of differences between Untreated-Doxo and RIPC-Doxo groups at 16 weeks was assessed by the Student *t*-test. (**B**) Wall thickening at 16 weeks in the doxorubicin-infused area (mean of the AHA segments 7 and 8). Grouped data are presented as mean±SE and individual data at the end of the study as a dotplot. N=10 animals per group. The significance of differences between Untreated-Doxo and RIPC-Doxo at 16 weeks was assessed by the Student *t*-test.

Improved LVEF after RIPC could be due to preserved contractile function in the doxorubicin-infused region or to a compensatory contractile effect in the remote area. To test these possibilities, we measured wall thickening as an index of contractility in the infused and remote LV regions (**Figure2.B**). RIPC was associated with a significant contractile improvement only in the infused region (23.8 % ± 15.2 and 6.59 ± 11.4 in RIPC-Doxo vs Untreated-Doxo; p = 0.01, **Figure 2.B**). Remote myocardium contractility did not differ significantly between groups (58.4 % ± 28.1 vs 45.4 ± 21.0 in RIPC-Doxo vs Untreated-Doxo; p = 0.26) (*Supplementary material online, Figure 2*).

### Remote ischemic preconditioning reduces myocardial fibrosis associated with anthracycline-induced cardiotoxicity

Myocardial tissue changes were evaluated in vivo by serial T1 mapping evaluations and ex-vivo in samples harvested at the end of the 16-week protocol. T1 relaxation times increased from baseline to week 16 in both treatment groups. In the RIPC-Doxo group, baseline T1 was 1150±58 ms and increased to 1300±109 ms at week 16 (p=0.03). In the Untreated-Doxo group, T1 increased from 1110±63 ms at baseline to 1410±108 ms at 16 weeks (p=0.01). The increase in T1 relaxation time was significantly smaller in the RIPC group (11.0±8.58% and 20.3±10.3% in RIPC-Doxo vs Untreated-Doxo; p = 0.04) (**Figure 3A**). Correlated with the CMR data, Sirius Red staining at the end of the study revealed a significantly smaller collagen area in the RIPC group (16.9 % ± 5.4 and 24.4 ± 9.6 in RIPC-Doxo vs Untreated-Doxo; p = 0.04) (**Figure 3B**).

**Figure 3:**
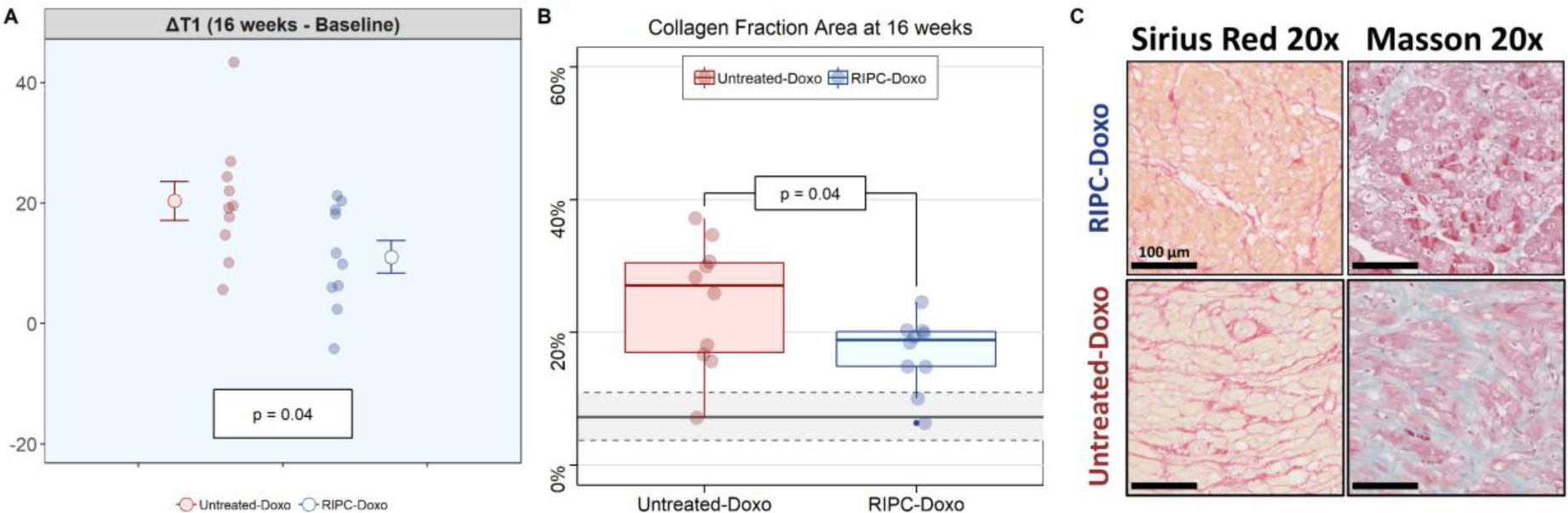
CMR and histology analysis of cardiac tissue. (**A**) Native T1 over time (left) and differential at the end of the study (16 weeks – baseline) (right). Grouped data are presented as mean±SE and individual data at the end of study as a dotplot. N=10 animals per group. The significance of differences between Untreated-Doxo and RIPC-Doxo groups at 16 weeks was assessed by the Student *t*-test. **B**) Collagen fraction area at 16 weeks. Data are presented grouped in a boxplot and ungrouped (individual data) as a dotplot. N=10 animals per group (9 in the Untreated-Doxo group for water content). The significance of differences between Untreated-Doxo and RIPC-Doxo at 16 weeks was assessed by the Student *t*-test. **C**) Representative histology images illustrating myocardial interstitial fibrosis (SR and Masson’s trichrome) at 16 weeks.

### Remote ischemic preconditioning attenuates mitochondrial fragmentation in end-stage anthracycline-induced cardiotoxicity

TEM evaluation of myocardial samples obtained at week 16 revealed massive mitochondrial fragmentation in non-RIPC pigs (median±IQR mitochondrial size 0.31±0.30 μm^2^ and 0.17±0.15 μm^2^ for healthy controls vs Untreated-Doxo pigs; p <0.001). Fragmentation was less severe in pigs undergoing RIPC before each doxorubicin injection (0.20±0.2 μm^2^ and 0.17±0.15 μm^2^ for RIPC-Doxo vs Untreated-Doxo; p < 0.001). Area distribution analysis confirmed that the size distribution of mitochondria in RIPC-Doxo pigs was between that of controls and animals (**Figure 4A-B**). In addition to the differences in mitochondria size, the Untreated-Doxo animals had elongated and aberrantly shape mitochondria, likely as a result of fragmentation; these, with massive loss of cristae and electrodense aggregates due to the subsequent cristae rupture (**Figure 4C**).

**Figure 4:**
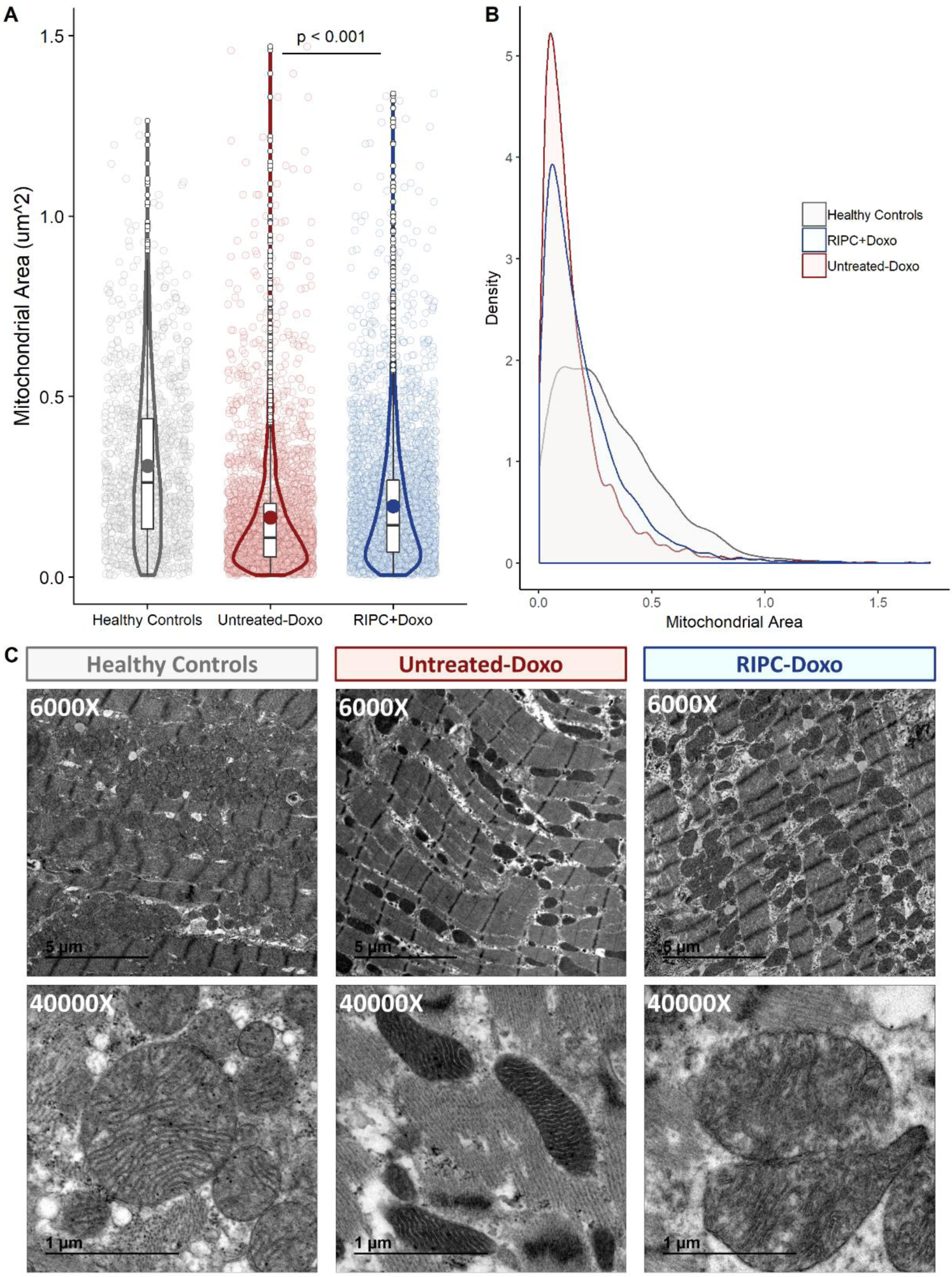
TEM analysis of myocardial mitochondria after 16 weeks of AIC (Study 1). (**A**) Individual mitochondrial area in the infused area in each animal group (dotplot) with its boxplot and violin plot related distribution. (**B**) Distribution of mitochondrial density vs mitochondrial area in each animal group. (**C**) Representative TEM images at 6000X and 40000X showing cardiac mitochondria in each animal group. N=10 animals (9 in the Untreated-Doxo group). Data were analyzed by linear regression nested according to treatment group and animal ID (Untreated-Doxo and RIPC-Doxo).

### Remote ischemic preconditioning prevents disruption of mitochondrial dynamics and dysregulated autophagy occurring early in the course of AIC

We next examined mitochondrial dynamics and other processes associated with cell damage during the subclinical stages of AIC, when cardiac function was still unaffected. For this, we included a new group of 10 pigs (Study 2) that received only three doxorubicin injections; as before, the pigs were first randomized to RIPC or no pretreatment (N=5 per arm). Pigs were sacrificed at week 6 (2 weeks after the third doxorubicin injection). CMR performed immediately before sacrifice revealed normal LVEF in both groups (57.7±2.08% and 55.3±5.16% in the RIPC-Doxo and Untreated-Doxo groups vs 64.1±6.24% in healthy controls). LV volumes and LV mass were also unaffected (*Supplementary material online, Table 2*).

Despite normal heart function, Untreated-Doxo pigs at this early stage of doxorubicin exposure had severe alterations in mitochondrial morphology, with a clear fragmented phenotype (median±IQR mitochondrial size = 0.27±0.34 μm^2^ in Untreated-Doxo pigs vs 0.308 ± 0.30 μm^2^ in healthy controls; p<0.001). This fragmented phenotype was not present in myocardial mitochondria from RIPC-Doxo pigs, and mitochondria in this group were significantly larger than in Untreated-Doxo pigs (0.34±0.31 μm^2^ vs 0.27±0.34 μm^2^ for RIPC-Doxo vs Untreated-Doxo; p<0.001) (**Figure 5A**). The area distribution curve for RIPC-Doxo pigs superimposed that of healthy controls, whereas Untreated-Doxo pigs showed a more fragmented distribution (**Figure 5B**). Irregularly shaped, small, and fragmented mitochondria surrounded by ribosomes were found in Untreated-Doxo pigs receiving only 3 doxorubicin injections (Study 2). The morphology of mitochondria from RIPC-Doxo pigs in Study 2 presented some disruption of cristae but conservation of shape and size (**Figure 5C**). Western blot analysis revealed that the mitochondrial fragmentation in Untreated-Doxo pigs was associated with a significant upregulation of the fission master regulator dynamin-related protein-1 (DRP1) (**Figure 6A and C**). Doxorubicin-induced DRP-1 upregulation was blocked RIPC (**Figure 6A and C**). mtDNA quantification confirmed the fragmented mitochondrial phenotype of Untreated-Doxo pigs, which had more mtDNA copies than RIPC-Doxo pigs and healthy controls (**Figure 6B**). Given the role of mitochondrial dysfunction in autophagy dysregulation and the impact on the latter on cardiac function, we next explored the impact of disrupted mitochondrial dynamics on the autophagy machinery. Expression of the autophagy-related proteins Beclin 1 and p62 was significantly higher in Untreated-Doxo pigs than in RIPC-Doxo pigs receiving doxorubicin and healthy controls (**Figure 6D-E**).

**Figure 5:**
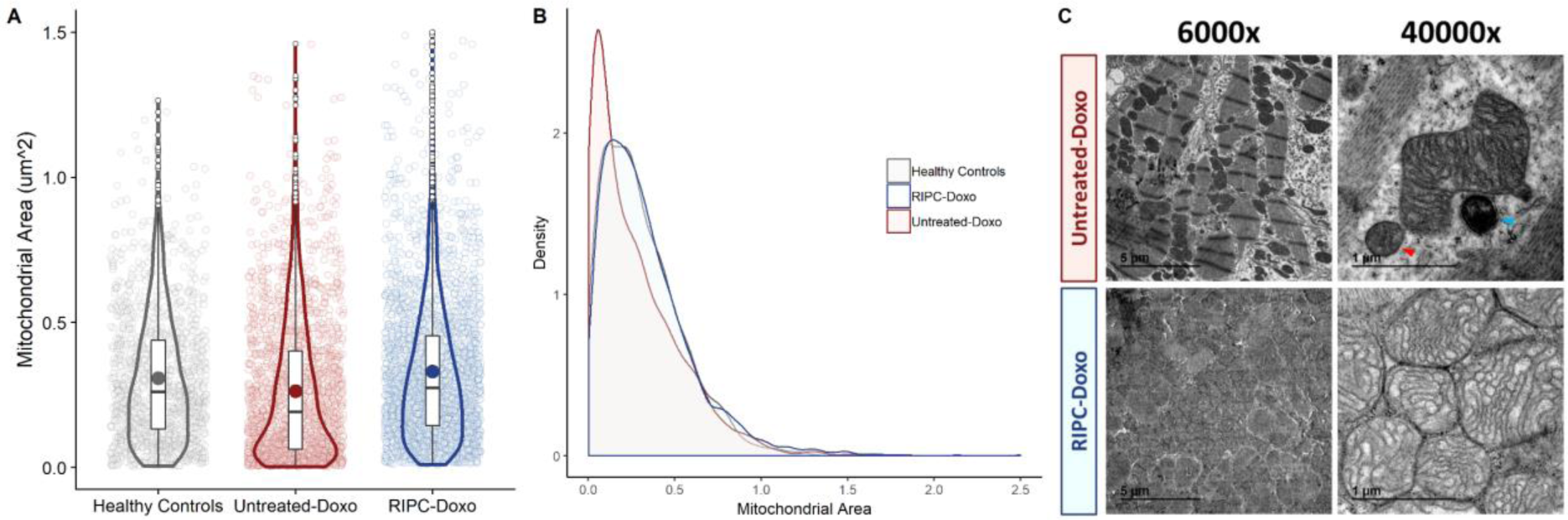
TEM analysis of myocardial mitochondria after 6 weeks of AIC (Study 2). (**A**) Individual mitochondrial area in the infused area in each animal group (dotplot) with its boxplot and violin plot related distribution. (**B**) Distribution of mitochondrial density vs mitochondrial area in each animal group. (**C**) Representative TEM images at 6000X and 40000X showing cardiac mitochondria in each animal group. N=5 animals. Data were analyzed by linear regression nested according to treatment group and animal ID (Untreated-Doxo and RIPC-Doxo). The red arrowhead indicates a fragmented mitochondria and the blue arrowhead a secondary lysosome.

**Figure 6:**
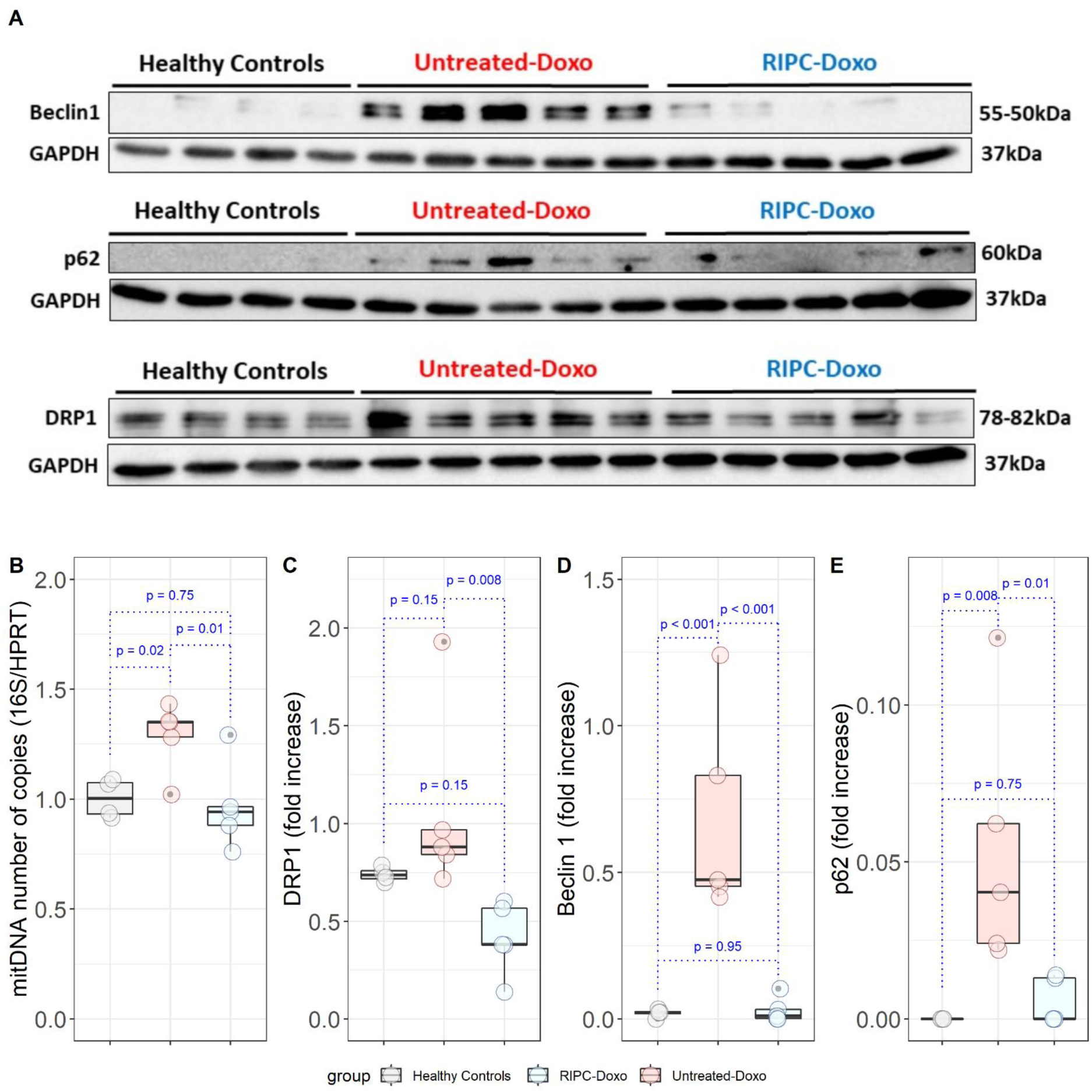
Myocardial mitochondria protein expression in the infused area after 6 weeks of AIC (Study 2). (**A**) Western blot analysis of the myocardial expression of Beclin1, p62, and DRP1 in all animals. (**B**) Myocardial mitochondrial DNA content. (**C-E**) Quantification of protein expression (fold-increased, normalized to GAPDH) for (**C**) DRP1, (**D**) Beclin 1, and (**E**) p62.

## Discussion

In this study, we tested the cardioprotective effect of RIPC before doxorubicin administration in a large-animal model of AIC. Long-term serial CMR evaluation showed that RIPC significantly ameliorates cardiac dysfunction associated with AIC. Hearts from RIPC-treated animals had significantly less myocardial fibrosis. Doxorubicin caused significant mitochondrial damage from the early stages after administration: at subclinical AIC stages (before overt cardiac dysfunction), doxorubicin induced a severe fragmented mitochondria phenotype and upregulated autophagy markers. RIPC applied before doxorubicin injections prevented the mitochondrial morphological abnormalities and the disruption of mitochondrial dynamics. These data identify RIPC as a potential cardioprotective strategy to prevent AIC and support the translation of this therapeutic strategy to the clinic.

Anthracyclines have been in clinical use for more than 50 years and, alone or in combination with other agents, remain the first line therapy for many cancer types. AIC is one of the most feared side effects of these efficacious chemotherapy agents. Most recent data show that up to 30% of patients treated with anthracyclines develop some degree of cardiotoxicity, and AIC manifests as significant cardiac dysfunction in up to 10% of patients^3,6^. Severe AIC is more frequent among especially vulnerable populations, such as the elderly. The unavoidable trade-off between cancer and chronic HF places a major burden on individuals and health care systems.

The current approach to AIC includes early detection by serial imaging and the use of non-specific HF therapies such as beta-blockers or ACE-inhibitors^4,15^. However, even when these are initiated early after AIC diagnosis, recovery cardiac function is usually incomplete^7^. Currently, the only FDA-approved drug for the prevention of AIC is dexrazoxane, an iron chelator that has been shown to reduce myocardial injury in a pediatric population^8,16^. However, dexrazoxane is not used in daily practice, and its efficacy for AIC prevention in adults is uncertain; more clinical trials are needed to clearly define its cardioprotective effect^17^. Moreover, most recent clinical trials with ACE inhibitors or betablockers showed inconsistent reductions in AIC^18–20^. There is therefore a clinical need to identify effective preventive therapies for AIC.

Remote ischemic conditioning has been extensively validated in experimental studies^10,21^ and early clinical trials^22^ as an intervention able to ameliorate cardiac damage associated with acute myocardial infarction (AMI). The protective effect of remote conditioning is stronger when it is applied before the index episode (PREconditioning); unfortunately, ischemia onset in AMI is an unpredictable event, and remote conditioning in this context is therefore only feasible during ongoing infarction (PERconditioning)^10^. However, in the largest clinical trial in AMI patients, the CONDI-2/ERIC-PPCI trial, remote ischemic perconditioning did not improve clinical outcomes^12^. Unlike AMI, anthracycline administration is a fully programmed event, and it is thus possible to schedule remote ischemic preconditioning. The positive results reported here and the ease of implementing this strategy in the clinic identify RIPC as a strong candidate for testing in future clinical trials. The safety of RIPC demonstrated in many trials in AMI add to the attraction of this strategy.

Ischemic conditioning has scarcely been tested in the context of AIC. Schjøtt et al^23^ tested ischemic preconditioning in isolated rat hearts before epirubicin administration; local ischemic preconditioning of explanted hearts was associated with an attenuation of epirubicin-induced cardiac dysfunction. More recently, Maulik et al^24^ tested ischemic preconditioning in cultured cardiomyocytes exposed to doxorubicin. Supporting the ex-vivo data^23^, their results showed that cardiomyocytes subjected to ischemic preconditioning were protected against doxorubicin-induced cell death^24^.

Importantly, ischemic preconditioning in these studies did not confer a competitive advantage to cancer cells exposed to doxorubicin, providing an additional safety benefit to this strategy. Very recently, Gertz et al^25^ published the first in vivo evidence of the cardioprotection afforded by RIPC in a mouse model of AIC. RIPC was associated with improved survival and attenuated cardiac fibrosis and autophagy. However, the anthracycline regime in that study did impact cardiac function, and thus the cardioprotective effect of RIPC was less clear^25^. In line with the Gertz et al study^25^, our results in pigs show that RIPC triggers a significant reduction in the cardiac expression of key autophagy factors.

Despite of the lack of experimental evidence from animal studies to support the ability of RIPC to prevent AIC, two small clinical trials are ongoing. Chung et al^26^ (NCT02471885) are testing a series of 4 RIPC cycles (5 minutes each) in the arm before each chemotherapy cycle in a population of 128 patients receiving anthracyclines for various types of cancer. In another trial (NCT03166813), Li et al are testing a different RIPC protocol (3 cycles of 5 minutes) applied either in the arm or in the leg before each chemotherapy cycle in a pediatric population (4-18 years old) receiving anthracyclines. Both trials have high-sensitivity cardiac troponin T (hs-cTnT) after the end of chemotherapy as the primary endpoint.

Our study provides the first demonstration that RIPC protects the heart against AIC in a large-animal model. Unlike the results from mice^25^, in our model, anthracyclines had a clear cardiocytotoxic effect, and we were able to detect differences in cardiac function and cardiac contractility between animals receiving RIPC or no pretreatment. We also found differences in T1 mapping values implicated in AIC^27^. In agreement with the results from mice^25^, at the end of the study protocol we found that RIPC-treated animals had significantly less fibrosis (collagen fraction) that animals receiving no pretreatment.

The mechanism by which RIPC exerts its cardioprotective effect against AIC is incompletely understood. However, there is firm evidence that mitochondrial damage plays a central role in AIC^2,28,29^. To investigate the mechanism of protection, we therefore explored mitochondrial dynamics (fusion/fission) and authophagy^30^. Our TEM analysis revealed that mitochondria from doxorubicin-injected pigs not undergoing RIPC were abnormally small, had irregular and aberrant morphology, and contained a hyper-dense matrix, confirming previous reports^28,31^. Morphological abnormalities were present very early in course of AIC (before overt cardiac dysfunction) and progressed to an extreme phenotype by the end of the 16-week protocol. RIPC prevented most of these morphological abnormalities from early in the course of AIC. Confirming these findings, protein expression of the mitochondrial fission master regulator DRP-1 was upregulated by doxorubicin treatment and normalized by RIPC, as was autophagy, evaluated early in the course of AIC by the expression of Beclin-1 and p62. Although autophagy is essentially protective, when dysregulated it can lead to cell death^32^. Mitochondrial dysfunction has emerged as essential trigger of autophagy. We propose that the severe mitochondrial dysfunction triggered by doxorubicin exposure induces a dysregulated autophagy in cardiomyocytes that leads to subsequent heart dysfunction. RIPC, by preserving mitochondrial integrity during AIC, halts this process and results in a long-term cardiac protection (**Summary Figure**).

In conclusion, this study provides evidence of the efficacy of RIPC in preventing AIC in a translational large-animal model. RIPC-mediated cardioprotection starts early in the course of AIC, in the subclinical phase, and has important downstream benefits on cardiac function. Our study supports the execution of adequately powered well-designed clinical trials to confirm this benefit in the clinical setting.

## Study limitations

The intracoronary route for doxorubicin injection used here does not mimic the clinical scenario and is a highly aggressive model. We^13^ and others^33,34^ have used this administration route in the past as a means of avoiding the myelosuppression associated with systemic doxorubicin injection, which is pronounced in pigs^35^. Nevertheless, the protective effect of RIPC in this extreme model strongly supports its translational potential. A further limitation in this study is the use of healthy juvenile pigs, which are free of the comorbidities frequent in cancer patients developing AIC, many of whom are elderly.

## Funding

This study is part of a project that has received funding from the European Research Council (ERC) under the European Union Horizon 2020 Research and Innovation Programme (ERC-Consolidator Grant agreement No. 819775 to B.I). The study was also partially funded by an ERA-CVD Joint Translational Call 2016 (funded through the Instituto de Salud Carlos III (ISCIII) and the European Regional Development Fund (ERDF), # AC16/00021) and by a Health Research Project from the ISCIII-FIS (# PI16/02110). Carlos Galán-Arriola and Rocío Villena-Gutiérrez are P-FIS fellows (Instituto de Salud Carlos III). This study forms part of a research agreement between the CNIC and Philips Healthcare. The CNIC is supported by the ISCIII, the Ministerio de Ciencia e Innovación, and the Pro-CNIC Foundation and is a Severo Ochoa Center of Excellence (MEIC award SEV-2015-0505).

## Acknowledgments

We thank Eugenio Fernández, Tamara Córdoba, Inés Sanz, Lorena Domínguez, Nuria Valladares, Antonio Benítez, Santiago Rodriguez-Colilla, and Rubén Mota for technical and veterinary support at the CNIC animal facility and farm. We also thank Marta Gavilán, Ángel Macías, and Braulio Pérez for technical support in CMR studies. Simon Bartlett (CNIC) provided English editing.

---

**Summary Figure:** Effect of RIPC on early and late stages of AIC.

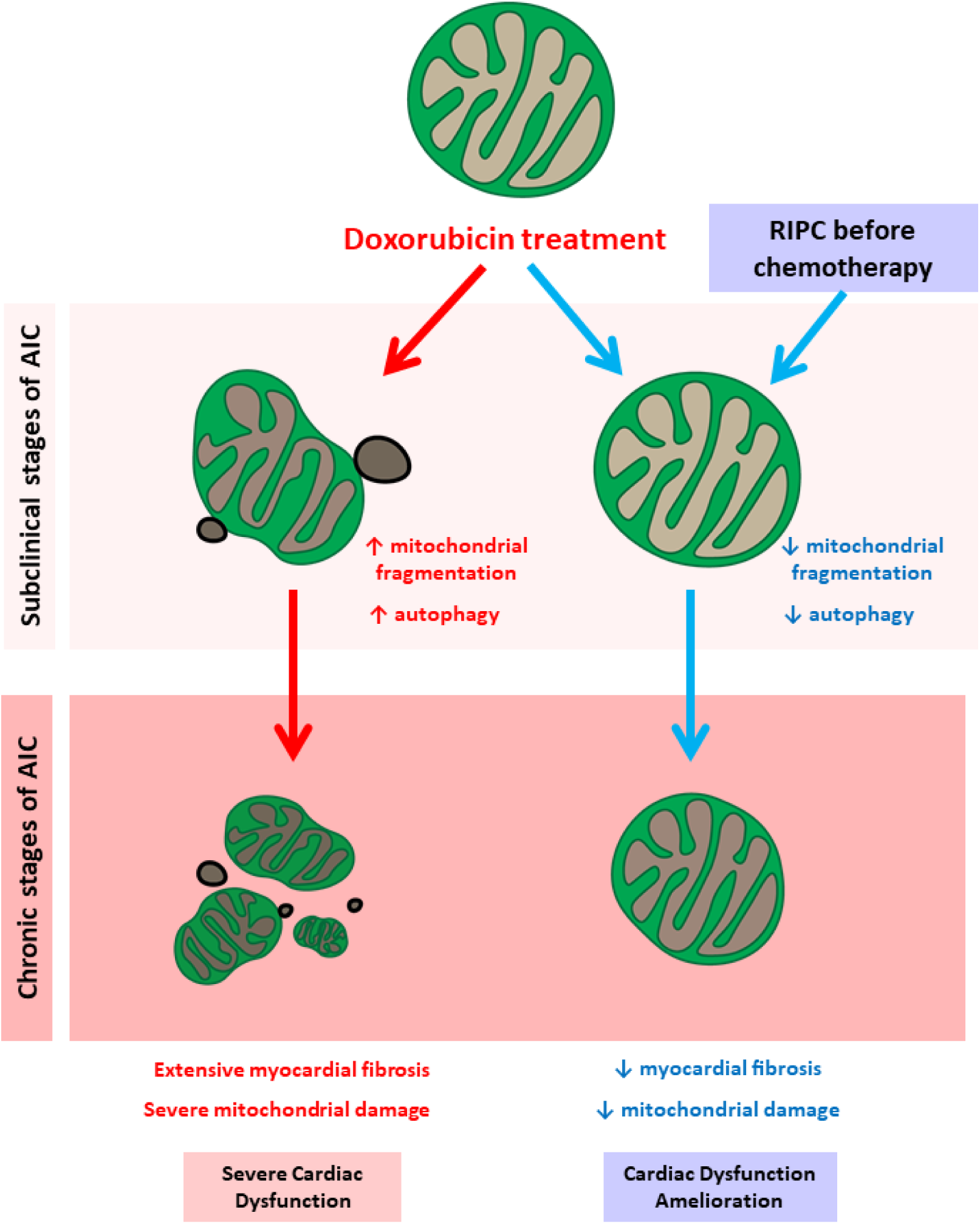

**Supplementary Figure 1.**
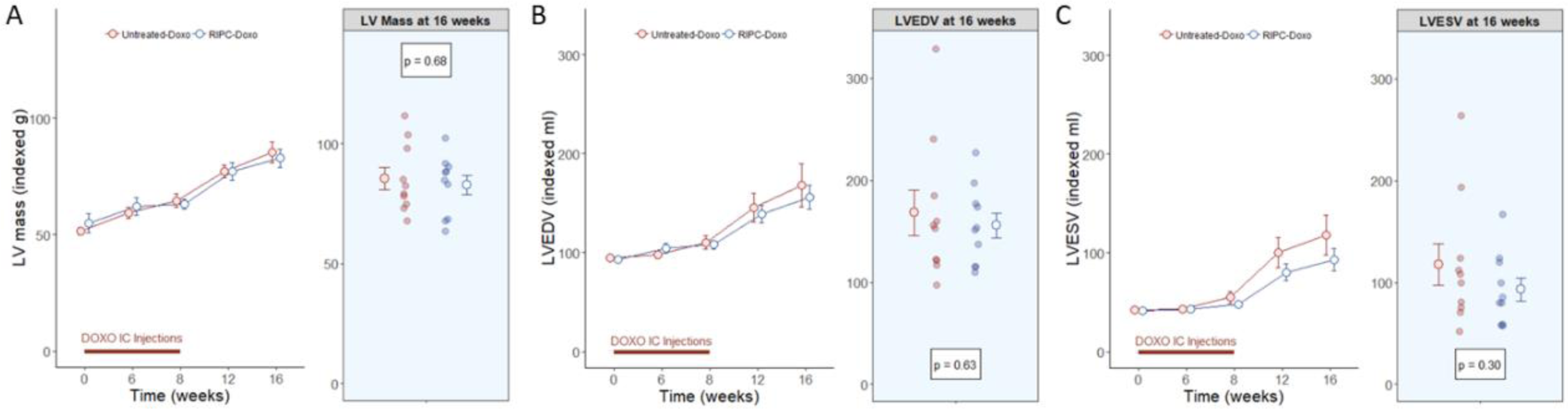
LV mass and volumes from experiment 1. Data by group, over time and at 16 weeks for (A) LV mass indexed, (B) LVEDV indexed and (C) LVESV indexed.

**Supplementary Figure 2.**
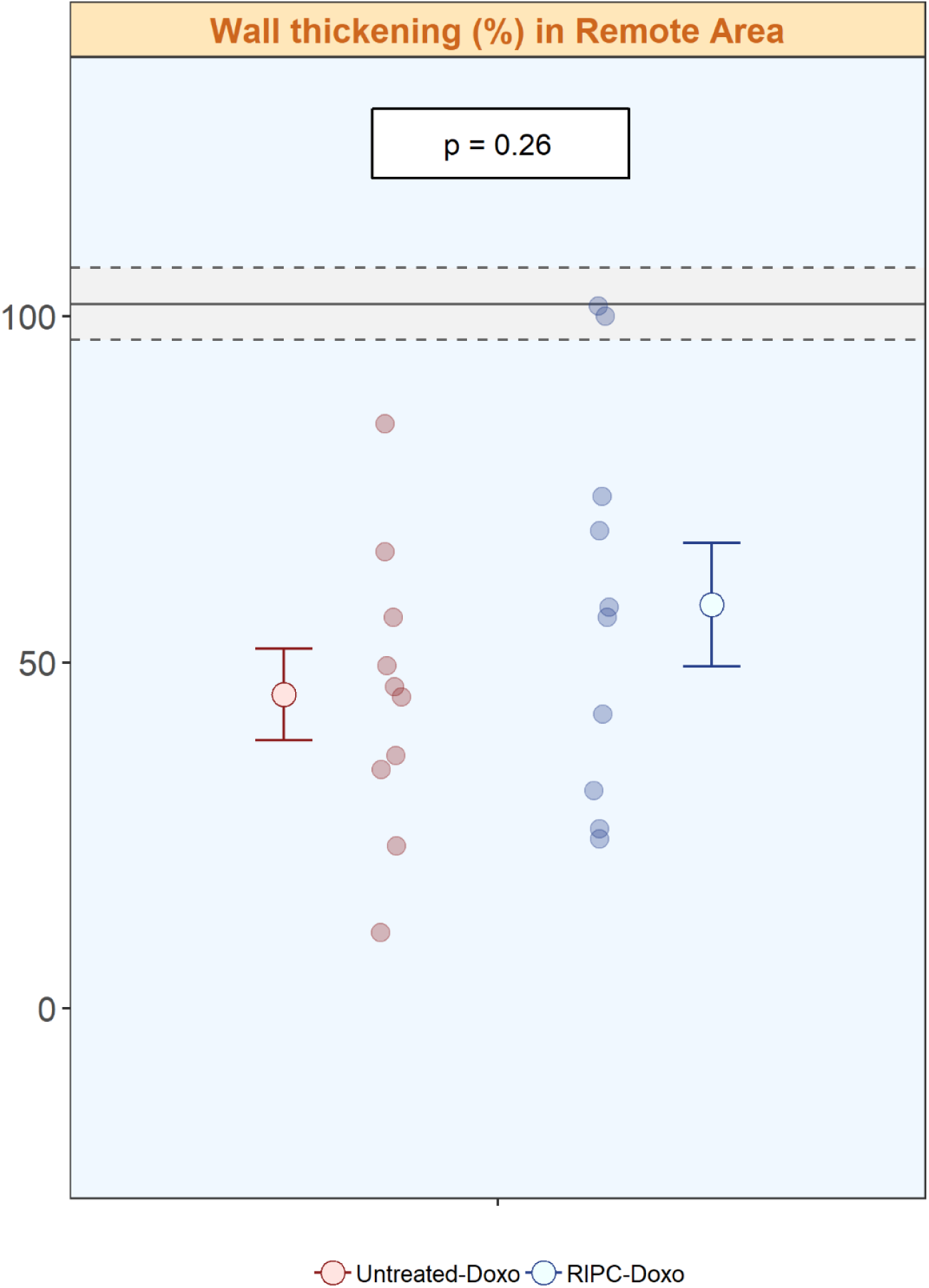
Cardiac contractility (wall thickening) at the end of the study for both groups in the remote area.

**Supplementary Table 1A.** CMR in-vivo data of the whole study 1 for all the time points and all the groups.

|  | 0 |  |  | 6 |  |  | 8 |  |  |
| --- | --- | --- | --- | --- | --- | --- | --- | --- | --- |
|  | Healthy Controls (n=10) | Untreated-Doxo (n=10) | RIPC-Doxo (n=10) | Healthy Controls (n=10) | Untreated-Doxo (n=10) | RIPC-Doxo (n=10) | Healthy Controls (n=10) | Untreated-Doxo (n=10) | RIPC-Doxo (n=10) |
| **LVEF (%)** |  |  |  |  |  |  |  |  |  |
| Mean (SD) | 56.9 (4.30) | 55.6 (5.64) | 55.5 (4.91) | 64.1 (6.24) | 55.7 (3.93) | 58.5 (5.24) | 61.8 (4.61) | 51.3 (6.41) | 55.5 (3.91) |
| Median [Min, Max] | 56.1 [50.9, 64.0] | 57.6 [45.8, 62.4] | 56.8 [48.3, 63.5] | 65.1 [52.7, 74.3] | 56.8 [48.8, 60.5] | 57.8 [50.3, 69.4] | 62.6 [55.0, 72.0] | 51.8 [39.0, 59.1] | 54.9 [50.1, 63.7] |
| **LV mass (mg indexed)** |  |  |  |  |  |  |  |  |  |
| Mean (SD) | 74.4 (9.28) | 51.5 (5.06) | 55.0 (13.3) | 51.3 (5.43) | 59.3 (7.59) | 62.1 (11.8) | 54.7 (4.55) | 64.5 (9.18) | 63.2 (6.26) |
| Median [Min, Max] | 71.8 [64.1, 97.4] | 52.7 [43.4, 59.3] | 51.9 [39.1, 87.6] | 50.2 [43.0, 60.2] | 60.4 [46.8, 69.9] | 58.7 [49.9, 86.1] | 54.1 [49.2, 64.0] | 63.9 [49.9, 78.7] | 62.8 [53.3, 75.6] |
| **LVEDV (ml indexed)** |  |  |  |  |  |  |  |  |  |
| Mean (SD) | 92.7 (10.3) | 94.8 (9.47) | 93.2 (10.9) | 92.3 (10.2) | 97.6 (10.5) | 105 (15.0) | 98.9 (9.82) | 110 (21.5) | 108 (13.4) |
| Median [Min, Max] | 91.3 [79.7, 111] | 96.2 [76.5, 106] | 96.1 [73.0, 109] | 88.0 [83.9, 113] | 98.2 [76.9, 115] | 106 [86.8, 131] | 100 [83.2, 112] | 103 [92.5, 152] | 105 [90.9, 127] |
| **LVESV (ml indexed)** |  |  |  |  |  |  |  |  |  |
| Mean (SD) | 40.2 (7.40) | 42.2 (7.16) | 41.6 (7.76) | 33.6 (9.32) | 43.3 (6.98) | 43.5 (8.22) | 37.8 (6.17) | 54.9 (18.3) | 48.2 (8.09) |
| Median [Min, Max] | 40.1 [30.8, 54.6] | 41.0 [31.6, 52.5] | 41.2 [26.7, 50.6] | 31.0 [22.3, 53.4] | 42.0 [31.7, 58.6] | 43.9 [26.6, 52.4] | 37.9 [28.6, 48.1] | 48.7 [40.0, 92.8] | 45.8 [36.2, 62.5] |
| **T2-GraSE (ms)** |  |  |  |  |  |  |  |  |  |
| Mean (SD) | 43.6 (1.72) | 43.2 (2.43) | 43.3 (1.79) | 44.6 (1.84) | 50.5 (2.60) | 52.2 (5.19) | 43.8 (2.60) | 55.2 (3.16) | 52.9 (4.41) |
| Median [Min, Max] | 43.6 [39.8, 45.7] | 43.5 [39.2, 48.1] | 43.0 [41.6, 46.6] | 43.6 [43.1, 48.6] | 50.3 [47.6, 56.6] | 51.8 [46.4, 65.2] | 43.5 [40.9, 49.5] | 55.1 [50.5, 61.4] | 52.7 [48.0, 61.5] |
| **T1 native (ms)** |  |  |  |  |  |  |  |  |  |
| Mean (SD) | 1160 (66.7) | 1110 (63.1) | 1150 (57.9) | 1220 (59.3) | 1120 (67.8) | 1160 (71.0) | 1150 (35.6) | 1210 (98.2) | 1220 (89.8) |
| Median [Min, Max] | 1130 [1100, 1300] | 1120 [1000, 1170] | 1150 [1070, 1220] | 1230 [1120, 1310] | 1120 [1000, 1260] | 1180 [1030, 1240] | 1140 [1120, 1230] | 1220 [996, 1320] | 1210 [1090, 1380] |
| **T1 post-contrast (ms)** |  |  |  |  |  |  |  |  |  |
| Mean (SD) | 807 (62.8) | 838 (65.6) | 809 (63.7) | 817 (214) | 778 (30.4) | 790 (43.1) | 776 (34.0) | 775 (61.8) | 742 (73.5) |
| Median [Min, Max] | 801 [684, 905] | 827 [772, 1010] | 826 [703, 886] | 756 [680, 1420] | 774 [722, 832] | 791 [733, 850] | 776 [726, 819] | 766 [656, 865] | 747 [562, 841] |
| **Extracellular Volume (%)** |  |  |  |  |  |  |  |  |  |
| Mean (SD) | 27.5 (4.64) | 24.8 (4.58) | 27.5 (2.14) | 30.3 (13.3) | 24.6 (3.64) | 27.8 (2.99) | 27.3 (2.59) | 29.4 (5.37) | 29.1 (8.18) |
| Median [Min, Max] | 27.0 [18.4, 35.1] | 23.6 [18.7, 31.3] | 27.4 [24.0, 31.3] | 34.6 [-6.86, 37.8] | 24.5 [19.0, 31.7] | 27.5 [23.5, 34.1] | 26.7 [24.2, 33.1] | 28.8 [18.8, 36.7] | 30.2 [7.63, 38.7] |

**Supplementary Table 1B.**
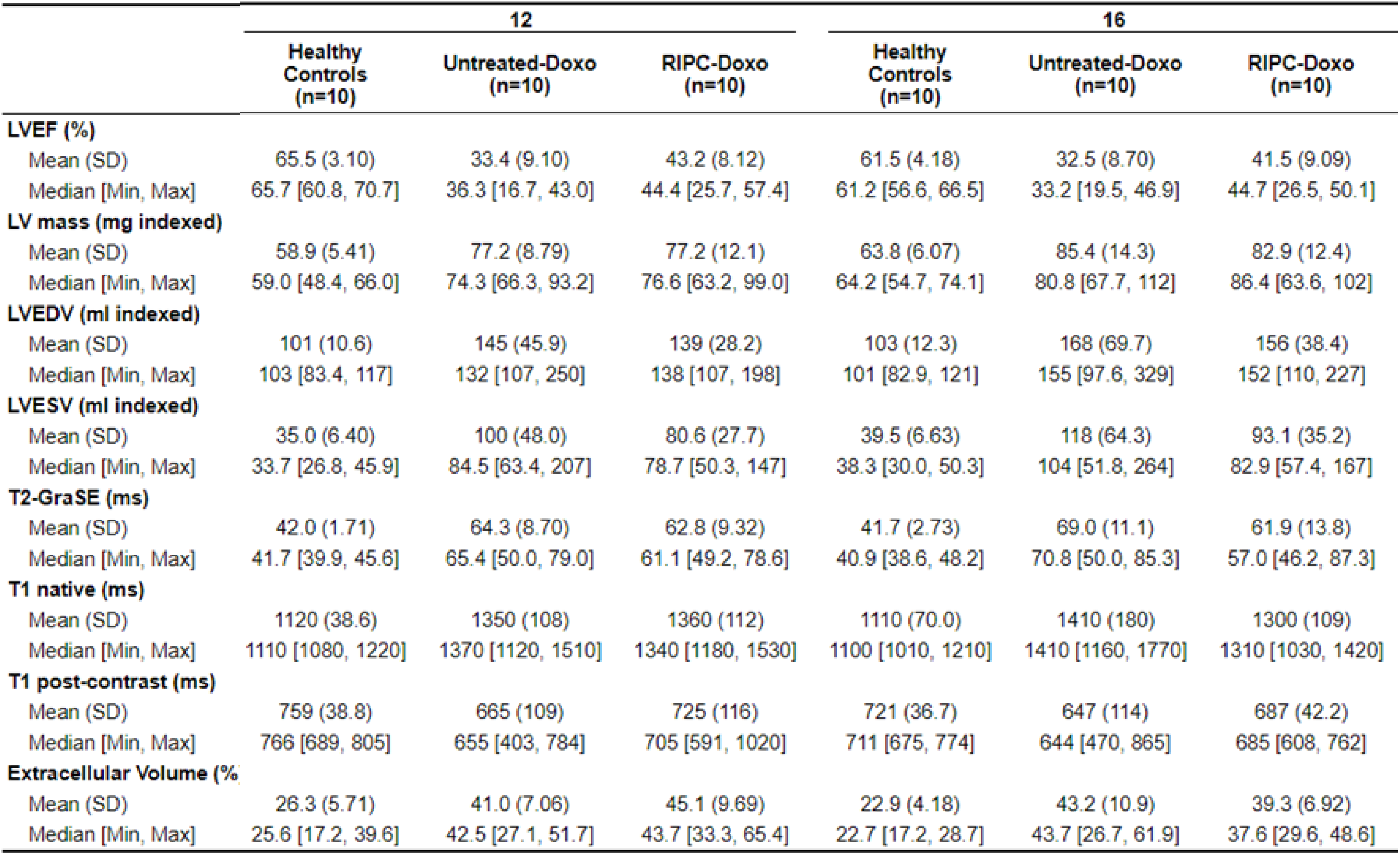
CMR in-vivo data of the whole study 1 for all the time points and all the groups

|  | 12 |  |  | 16 |  |  |
| --- | --- | --- | --- | --- | --- | --- |
|  | Healthy Controls (n=10) | Untreated-Doxo (n=10) | RIPC-Doxo (n=10) | Healthy Controls (n=10) | Untreated-Doxo (n=10) | RIPC-Doxo (n=10) |
| **LVEF (%)** |  |  |  |  |  |  |
| Mean (SD) | 65.5 (3.10) | 33.4 (9.10) | 43.2 (8.12) | 61.5 (4.18) | 32.5 (8.70) | 41.5 (9.09) |
| Median [Min, Max] | 65.7 [60.8, 70.7] | 36.3 [16.7, 43.0] | 44.4 [25.7, 57.4] | 61.2 [56.6, 66.5] | 33.2 [19.5, 46.9] | 44.7 [26.5, 50.1] |
| **LV mass (mg indexed)** |  |  |  |  |  |  |
| Mean (SD) | 58.9 (5.41) | 77.2 (8.79) | 77.2 (12.1) | 63.8 (6.07) | 85.4 (14.3) | 82.9 (12.4) |
| Median [Min, Max] | 59.0 [48.4, 66.0] | 74.3 [66.3, 93.2] | 76.6 [63.2, 99.0] | 64.2 [54.7, 74.1] | 80.8 [67.7, 112] | 86.4 [63.6, 102] |
| **LVEDV (ml indexed)** |  |  |  |  |  |  |
| Mean (SD) | 101 (10.6) | 145 (45.9) | 139 (28.2) | 103 (12.3) | 168 (69.7) | 156 (38.4) |
| Median [Min, Max] | 103 [83.4, 117] | 132 [107, 250] | 138 [107, 198] | 101 [82.9, 121] | 155 [97.6, 329] | 152 [110, 227] |
| **LVESV (ml indexed)** |  |  |  |  |  |  |
| Mean (SD) | 35.0 (6.40) | 100 (48.0) | 80.6 (27.7) | 39.5 (6.63) | 118 (64.3) | 93.1 (35.2) |
| Median [Min, Max] | 33.7 [26.8, 45.9] | 84.5 [63.4, 207] | 78.7 [50.3, 147] | 38.3 [30.0, 50.3] | 104 [51.8, 264] | 82.9 [57.4, 167] |
| **T2-GraSE (ms)** |  |  |  |  |  |  |
| Mean (SD) | 42.0 (1.71) | 64.3 (8.70) | 62.8 (9.32) | 41.7 (2.73) | 69.0 (11.1) | 61.9 (13.8) |
| Median [Min, Max] | 41.7 [39.9, 45.6] | 65.4 [50.0, 79.0] | 61.1 [49.2, 78.6] | 40.9 [38.6, 48.2] | 70.8 [50.0, 85.3] | 57.0 [46.2, 87.3] |
| **T1 native (ms)** |  |  |  |  |  |  |
| Mean (SD) | 1120 (38.6) | 1350 (108) | 1360 (112) | 1110 (70.0) | 1410 (180) | 1300 (109) |
| Median [Min, Max] | 1110 [1080, 1220] | 1370 [1120, 1510] | 1340 [1180, 1530] | 1100 [1010, 1210] | 1410 [1160, 1770] | 1310 [1030, 1420] |
| **T1 post-contrast (ms)** |  |  |  |  |  |  |
| Mean (SD) | 759 (38.8) | 665 (109) | 725 (116) | 721 (36.7) | 647 (114) | 687 (42.2) |
| Median [Min, Max] | 766 [689, 805] | 655 [403, 784] | 705 [591, 1020] | 711 [675, 774] | 644 [470, 865] | 685 [608, 762] |
| **Extracellular Volume (%)** |  |  |  |  |  |  |
| Mean (SD) | 26.3 (5.71) | 41.0 (7.06) | 45.1 (9.69) | 22.9 (4.18) | 43.2 (10.9) | 39.3 (6.92) |
| Median [Min, Max] | 25.6 [17.2, 39.6] | 42.5 [27.1, 51.7] | 43.7 [33.3, 65.4] | 22.7 [17.2, 28.7] | 43.7 [26.7, 61.9] | 37.6 [29.6, 48.6] |

**Supplementary Table 2.**
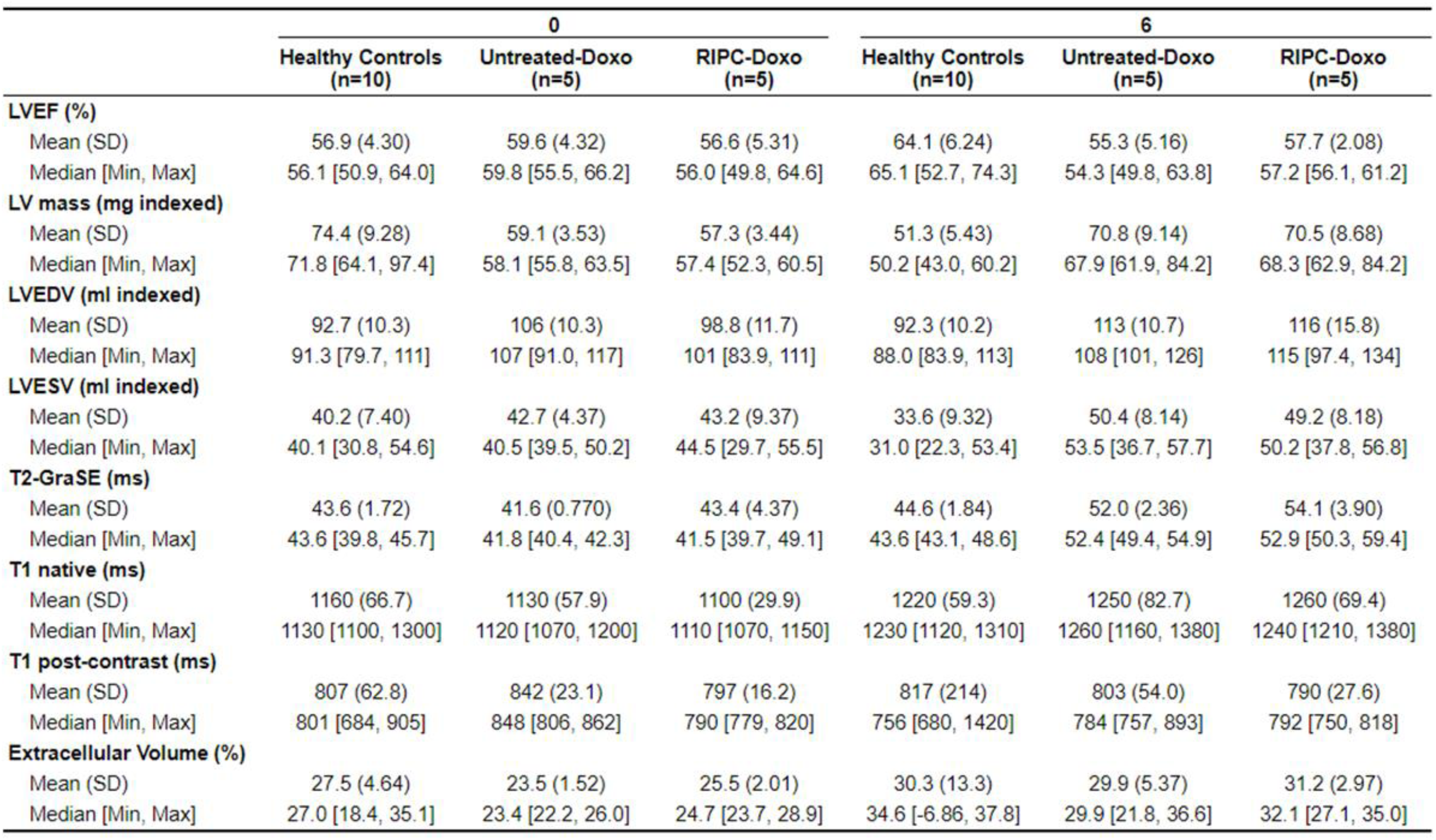
CMR in-vivo data of the whole study 2 for all the time points and all the groups (healthy controls at 0 and 6 weeks from study 1 are also included for comparison).

